# Tissue specific requirements for ECSIT in mitochondrial complex I assembly

**DOI:** 10.1101/2021.04.01.438015

**Authors:** Thomas L Nicol, Sara Falcone, Andrew Blease, Pratik Vikhe, Gabriele Civiletto, Saleh Salman Omairi, Carlo Viscomi, Ketan Patel, Paul K Potter

**Affiliations:** MRC Harwell Institute, Mammalian Genetics Unit, Harwell Campus, Oxfordshire, UK; BHF Centre of Research Excellence, Division of Cardiovascular Medicine, Radcliffe Department of Medicine, John Radcliffe Hospital, University of Oxford, Oxford, UK; Centre for Cellular and Molecular Physiology, University of Oxford, Oxford, UK; MRC Mitochondrial Biology Unit, University of Cambridge, Cambridge, UK; School of Biological Sciences, University of Reading, Reading, UK; Department Biological and Medical Sciences, Faculty of Health and Life Sciences, Oxford Brookes University, UK

**Keywords:** complex I, *Ecsit*, hypertrophic cardiomyopathy, mitochondria

## Abstract

Here we describe a mutation in the mitochondrial complex I assembly factor (Evolutionarily conserved signalling intermediate in Toll pathway) ECSIT which reveals tissue specific requirements for this factor in complex I assembly. Mitochondrial complex I assembly is a multi-step process dependant on assembly factors that organise and arrange the individual subunits, allowing for their incorporation into the complete enzyme complex. We have identified an ENU induced mutation in ECSIT (N209I) that exhibits a profound effect on complex I assembly only in heart tissue resulting in hypertrophic cardiomyopathy in the absence of other phenotypes. Mitochondrial function was reduced by 98% in mitochondria isolated from cardiac tissue but mitochondria from other tissues such as skeletal muscle, brain, liver, and kidney were unaffected. This data suggests the mechanisms underlying complex I assembly are tissue specific and has implications in understanding the pathogenesis of cardiomyopathy.

## Introduction

Given the size, complexity and contributions from the mitochondrial and nuclear genomes, it is unsurprising that the assembly of mitochondrial complex I is an intricate process which we are only beginning to understand. Complex I consists of 45 subunits, 7 of which are encoded by the mitochondrial DNA and the remaining in the nucleus. Involved in its assembly are at least 14 assembly factors which are not thought to comprise part of the final structure but are essential in the intervening steps between isolated proteins and functional complex (Guerrero-Castillo *et al*, 2016; Mimaki *et al*, 2012).

The assembly process proceeds in a stepwise fashion, with individual building blocks or sub-assemblies forming first, before joining to form structural or functional portions of the complex and ultimately the complete complex. There are 4 main modules that must be assembled for full function: N, the NADH binding domain; Q, the quinone binding domain; P_p_ and P_D_, the proximal and distal portions of the membrane arm involved in proton pumping (Sanchez-Caballero *et al*, 2016a).

Amongst the assembly factors involved in complex I assembly is the evolutionarily conserved signalling intermediate in toll pathway (ECSIT). In humans ECSIT is a 431 amino acid adapter protein with 2 identifiable isoforms (50/33kDa) and a third potential isoform based on splice prediction (24kDa) (Kopp *et al*, 1999; Xiao *et al*, 2003).

Human ECSIT protein has 3 recognisable domains in the full length protein, an N-terminal mitochondrial targeting sequence (amino acids 1-48), a highly ordered pentatricopeptide repeat region (PPR) (amino acids 90-266) and a less ordered C terminal domain that shows some 3D resemblance to pleckstrin homology domains (amino acids 275-380) (Giachin *et al*, 2016). Mouse ECSIT protein maintains roughly 73% sequence homology to the human protein, with the same mitochondrial targeting sequence seen in the human protein (amino acids 1-48) (results according to Phyre2 web server) (Kelley *et al*, 2015).

ECSIT was first identified as interacting with the proteins TRAF6 (tumour necrosis factor (TNF) receptor associated factor 6) and MEKK-1/MAP3K1 (ERK kinase kinase-1/Mitogen–activated protein kinase) in the Toll/IL-1 pathway. It has been shown that ECSIT binds to the multi-adapter protein TRAF6 and allows for the phosphorylation of MEKK1 (MAP3K1) into an active state. This phosphorylation event leads to activation of NF-κB and promotion of the innate immune response (Kopp *et al*., 1999).

During complex I assembly, ECSIT interacts with NDUFAF1 and complex I, and has previously been identified in large complex I assembly complexes of approximately 500, 600 and 850 kDa along with NDUFAF1. siRNA knock down of ECSIT results in loss of complex I protein levels and enzymatic activity (Giachin *et al*., 2016; Vogel *et al*, 2007). ACAD9 is another complex I assembly factor that interacts with both ECSIT and NDUFAF1 and knock down of ACAD9 leads to a decrease in NDUFAF1 and ECSIT protein levels and a reduction in functional complex I levels (Nouws *et al*, 2010). Together these 3 proteins form part of the mitochondrial complex I assembly complex (MCIA) along with TMEM126B and TIMMDC1 (Giachin *et al*., 2016; Heide *et al*, 2012). ACAD9 forms a homodimer which acts as the scaffold for the interaction of ECSIT and NDUFAF1, bringing the 3 proteins together. This trimer then interacts with the membrane bound proteins TMEM126B and TIMMDC1 before acting as part of the complex I assembly process (Giachin *et al*., 2016).

Patients with complex I deficiencies display a wide variety of phenotypes varying from syndromes, such as Leigh syndrome (Loeffen *et al*, 1998; Rahman *et al*, 1996) and MELAS syndrome (Pavlakis *et al*, 1984; Sproule & Kaufmann, 2008) which may arise in early childhood, to later onset or milder conditions. Amongst these, many other clinical features have been reported, including cases of exercise intolerance, renal tubular acidosis, lactic acidosis, cardiomyopathy and encephalopathy due to mutations in complex I subunits (NDUFV2 and NDUFS2) (Benit *et al*, 2003; Loeffen *et al*, 2001) or assembly factors (ACAD9, NDUFAF1 and TMEM126B) (Alston *et al*, 2016; Fassone *et al*, 2011; Nouws *et al*., 2010; Sanchez-Caballero *et al*, 2016b). Mutations in complex I assembly factors results in a range of disease phenotypes (Ghezzi & Zeviani, 2018) which may suggest variability in the sensitivity of different tissues to deficiencies in complex I activity.

Here we describe a novel, ENU induced mutation (Potter *et al*, 2016) in the complex I assembly factor ECSIT which results in a profound cardiomyopathy phenotype with other tissues apparently unaffected. We demonstrate that this phenotype arises from an impairment of complex I assembly in cardiac tissue, which is maintained at or close to normal levels in other tissues tested. Furthermore, the evidence indicates that ECSIT has a tissue specific role in the complex I assembly process and could shed light on the differences in phenotypes observed in complex I deficient patients.

## Results

### Phenotype identification and genetic characterisation

As part of a phenotype-driven screen to identify mutations resulting in age-related and chronic disease we identified a group of related mice (Potter *et al*., 2016) exhibiting various signs of ill health (sudden weight loss, hunched appearance, piloerect coat, inactivity) or that died unexpectedly. *Post mortem* analysis revealed enlarged hearts in these mice and histological analysis showed characteristic signs of hypertrophic cardiomyopathy (HCM): enlargement and disorganisation of the cardiomyocytes and the presence of vacuolation (**Fig EV1**).

The causative mutation was mapped to a 46Mb region at the proximal end of chromosome 9 (**Fig 1 A**). Whole genome sequencing (WGS) identified a single coding variant within the mapping region, an A to T transversion at nucleotide 916 of the gene *Ecsit*. This mutation was validated by Sanger sequencing (**Fig 1 B**) and shown to lie in the predicted penta-tricopeptide repeat (PPR) motif of the mouse ECSIT protein (**Fig 1 C**) (protein domains predicted by Phyre 2 webserver (Kelley *et al*., 2015)). *Ecsit^N209I/+^* mice were crossed to Ecsit^+/-^ animals to produce the compound heterozygotes and other intermediate genotypes. Heart weights from the 4 genotypes produced (**Fig 1 D-E**) demonstrate that the phenotype is only present in compound heterozygotes, thus confirming that the mutation in *Ecsit* is the causative allele.

**Fig 1:**
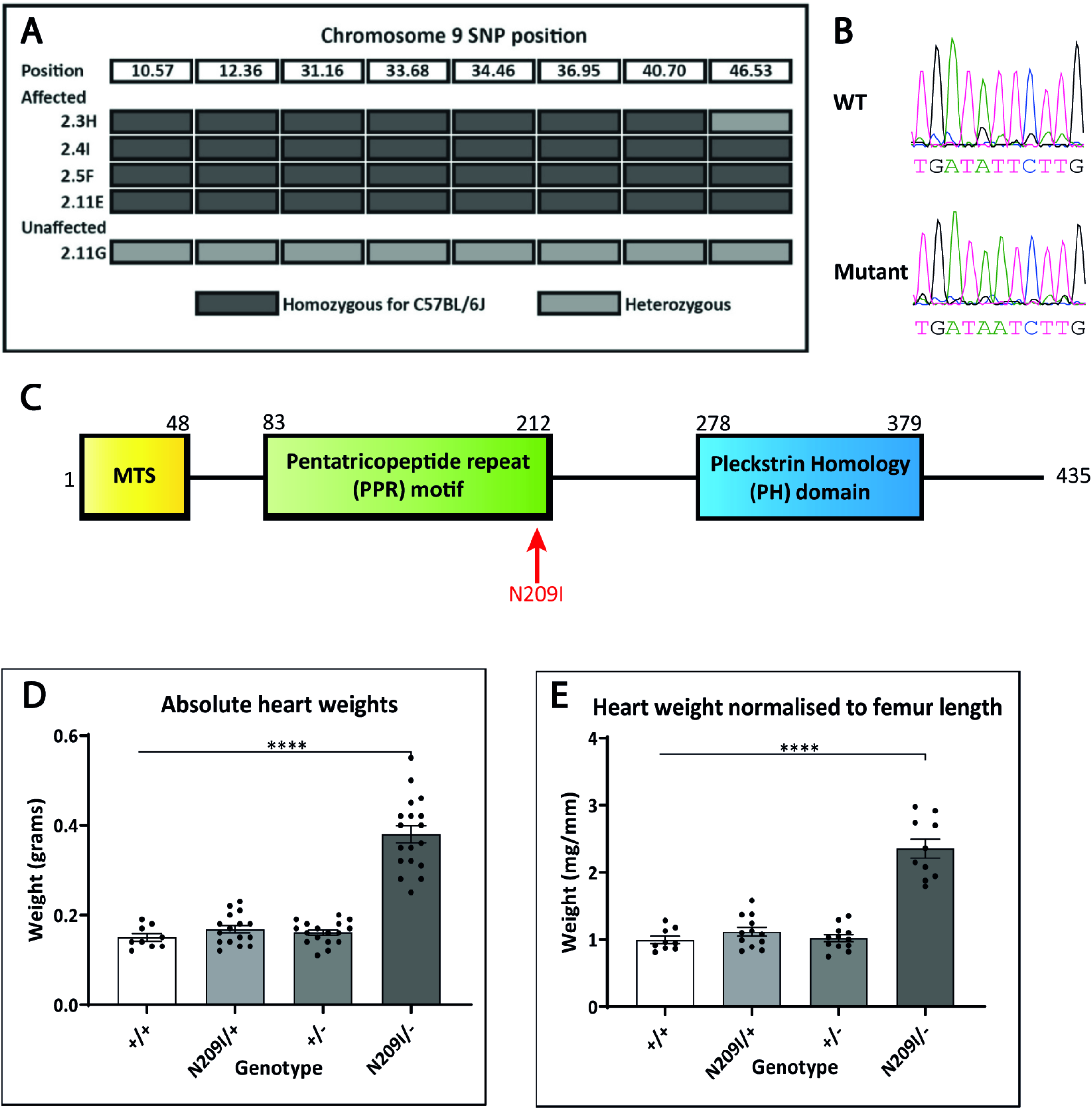
(A) Chromosome 9 mapping location identified by SNP mapping. All affected animals are homozygous for SNPs derived from the C57BL/6J founder between the proximal end and 46.53Mb. (B) Sanger sequencing confirmation of an A to T transversion at position 916 of *Ecsit*. (C) heart weights from wild type, heterozygous N209I mutant, heterozygous knockout, and compound heterozygote animals confirm that loss of ECSIT protein function is causative of the hypertrophy phenotype observed. (D) Normalised heart weights from wild type, heterozygous N209I mutant, heterozygous knockout, and compound heterozygote animals. Mean ± SEM, ****p<0.0001.

### Cardiac Phenotyping

The mutant line was backcrossed to C3H.Pde6b+ for 5 generations to produce an incipient congenic line used for all further phenotype characterization. Hearts from wild-type and *Ecsit^N209I/N209I^* animals were collected as a time course (birth, 1, 2, 4, 6, 8 and 12 weeks, 12-week data is shown (**Fig 2 A**)), to determine the onset of disease and characterize the progression. Signs of HCM (vacuolation, mineralization, myocyte disorganisation) were present from 6 weeks of age with progression (myocyte hypertrophy) clear at 8 and 12 weeks of age. In addition, tissue weights (**Fig 2 B**), both in absolute values and normalised to body weight or femur length, demonstrated a significant increase in heart weight of *Ecsit^N209I/N209I^* animals compared to controls at 12 weeks of age. Lung weights also showed a significant increase in *Ecsit^N209I/N209I^* animals suggestive of lung congestion resulting from left ventricular hypertrophy, conversely, liver weights were not increased, suggesting that the right heart was not congested.

**Fig 2:**
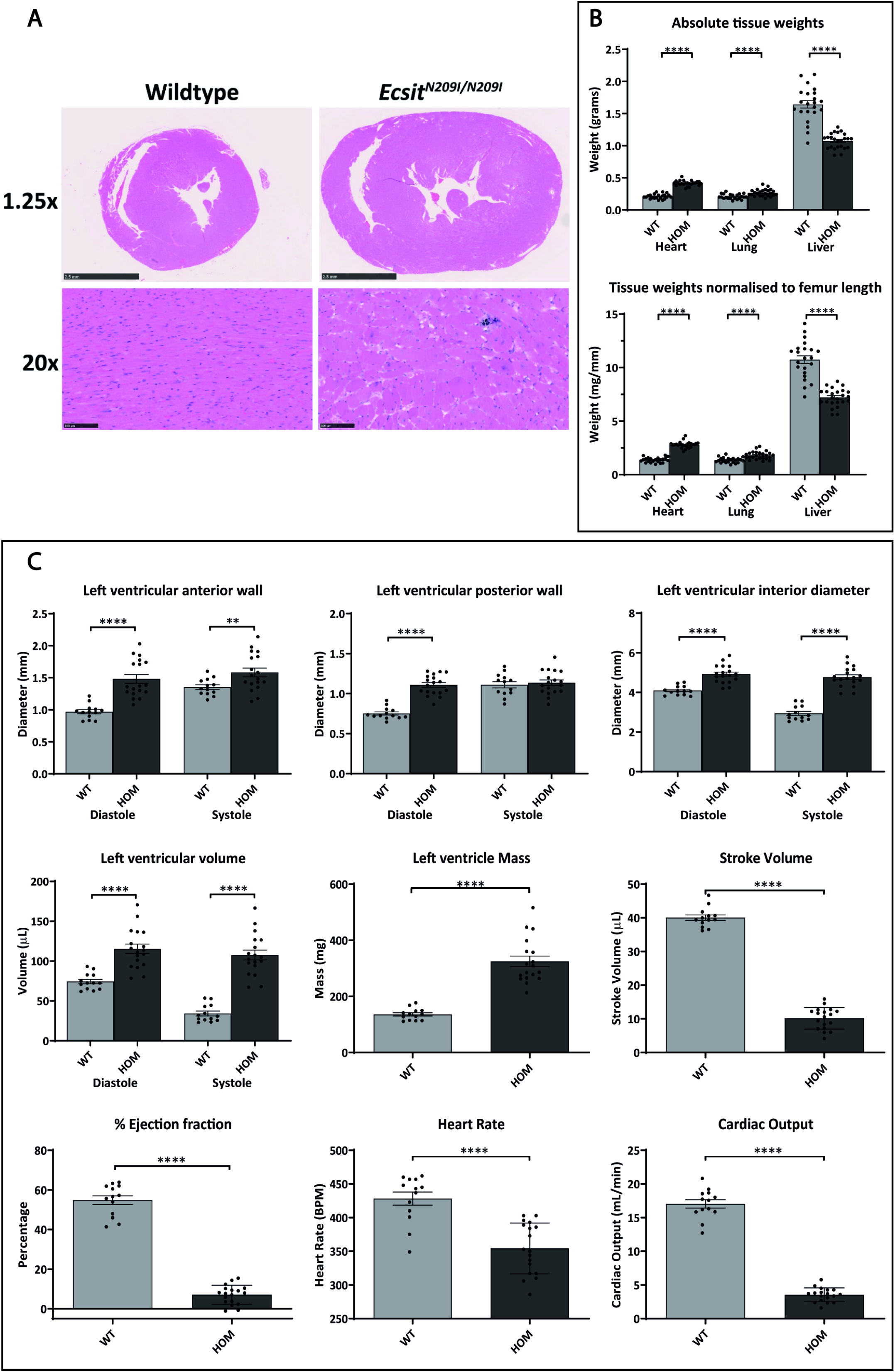
(A) Histology of wild-type and *Ecsit^N209I/N209I^* hearts demonstrating overall size increase as well as disorganised, enlarged and vacuolated cardiomyocytes, indicative of HCM. (B) Absolute and normalised (to femur length) weights of heart, lung and liver in backcrossed animals demonstrating phenotype is still present and that congestion is primarily of the left heart. (C) M-mode measurements of echocardiography showing significant enlargement of anterior and posterior wall as well as an increase in ventricular volume and mass. Furthermore, key measurements of cardiac function, stroke volume, ejection fraction and cardiac output are significantly reduced. Mean ± SEM, **p<0.01, ****p<0.0001.

Echocardiography was performed on animals at 12 weeks of age (**Fig 2 C**) demonstrating a thickening of the left ventricular anterior and posterior walls coupled with an overall increase of left ventricular mass and volume. Stroke volume, ejection fraction and cardiac output all demonstrate a profound reduction in *Ecsit^N209I/N209I^* animals in comparison to wild type. Taken together these data confirm the presence of HCM and suggest that there may also be some dilation of the left ventricle.

Further phenotyping by Echo-MRI revealed a reduction in total body weight, fat mass and lean mass in *Ecsit^209I/N2091^* animals from 14 weeks of age (**FigFig EV2**). No difference was seen in ECG parameters (data not shown) or in muscle fibre types of the soleus and extensor digitorum longus (EDL) muscles of the hind limb (**FigFig EV3 A-B**). Muscle fibre analysis did reveal a significant reduction in cross sectional area of both the soleus and EDL in *Ecsit^N209I/N209I^* animals in comparison to wild type (**FigFig EV3 C-D**), although it is unclear if this is a result of the general reduction in size of the mutant animals (**Fig 2 A-B**) or a true muscle phenotype.

Taken together this data confirms an HCM phenotype in the *Ecsit^N209I/N209I^* animals in comparison to wild type. This phenotype is present in the absence of other profound muscular phenotypes although may be involved in the development of overall body mass and fat reserves.

### Identification of pathogenic pathway underlying the hypertrophic cardiomyopathy

ECSIT has been shown previously to be involved in both the Toll-like Receptor (TLR) response (Kopp *et al*., 1999; Wi *et al*, 2014) and in the assembly of complex I of the mitochondrial electron transport chain (Giachin *et al*., 2016; Vogel *et al*., 2007), both of which could be considered as causes for the development of a HCM phenotype. To determine which pathway was the underlying driver for the development of this phenotype, characterisation of both the TLR response and mitochondrial electron transport assembly was undertaken.

Mitochondrial electron transport chain proteins NDUFB8 (CI), SDHA (CII), UQCRC2 (CIII), MTCO1 (CIV) and ATP5A (CV) were assessed in whole heart lysate from male and female animals at 16 weeks of age by western blot. A 98% reduction (p=0.0046) was present in complex I protein levels in the heart (**Fig 3 A**). No differences were observed in any other electron transport chain proteins. In addition, complex I protein levels were also reduced in brain (~30%), kidney (~30%) and liver (~60%) (**FigFig EV4**).

**Fig 3:**
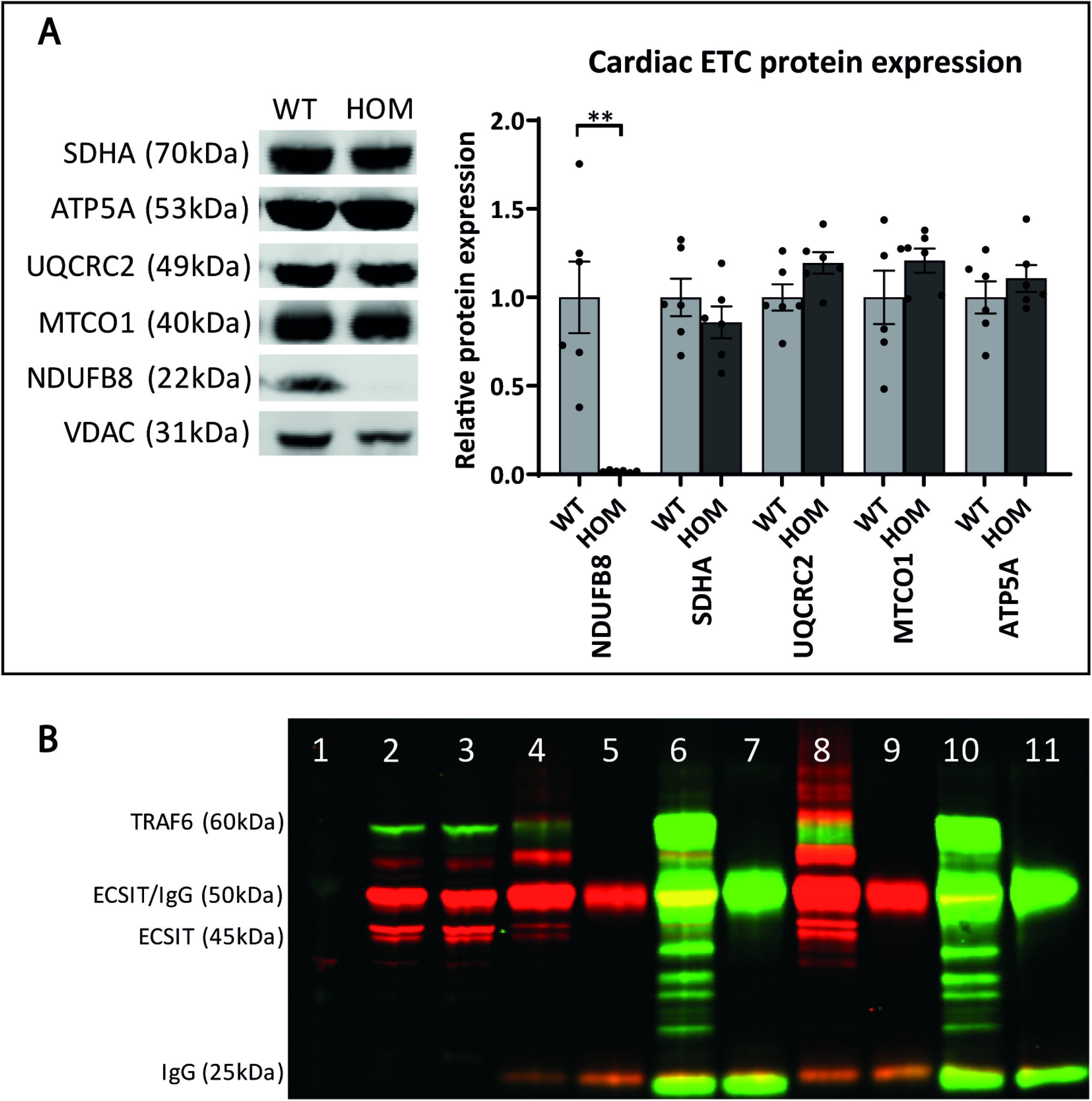
(A) Representative cardiac electron transport chain protein levels for each complex (I – NDUFB8, II – SDHA, III – UQCRC2, IV – MTCO1, V – ATP5A). (B) Immunoprecipitation of wild-type and mutant ECSIT (His tagged) (45 and 50kDa) with full length wild-type TRAF6 (Myc tagged) (60kDa) (green – Myc) and rabbit (red – His) 1. Untransfected input lysate, 2. Wild-type ECSIT(His) + wild-type TRAF6(Myc) input lysate 3. ECSIT N209I(His) + wild-type TRAF6(Myc) input lysate, 4. Wild-type ECSIT(His) + wild-type TRAF6(Myc) anti-His immunoprecipitation, 5. Empty AC-His vector + wild-type TRAF6(Myc) anti-His immunoprecipitation, 6. Wild-type ECSIT(His) + wild-type TRAF6(Myc) anti-Myc immunoprecipitation, 7. Wild-type ECSIT(His) + empty entry(Myc) vector anti-Myc immunoprecipitation, 8. N209I ECSIT(His) + wild-type TRAF6(Myc) anti-His immunoprecipitation, 9. Empty AC-His vector + wild-type TRAF6(Myc) anti-His immunoprecipitation, 10. N209I ECSIT(His) + wild-type TRAF6(Myc) anti-Myc immunoprecipitation, 11. N209I ECSIT + empty entry(Myc) vector anti-Myc immunoprecipitation. Mean ± SEM, **p<0.01.

To assess the role of the TLR response in development of the cardiomyopathy phenotype FACS analysis on whole blood from 12-week-old male and female animals was performed to determine the constituents of the total leukocyte fraction. Results demonstrate a small reduction in the percentage of circulating lymphocytes and macrophages with no differences in monocyte or neutrophil levels (**FigFig EV5 A**).

In addition, bone marrow derived macrophages (BMDMs) were cultured from the bone marrow of 12-week-old *Ecsit^N209I/N209I^* and *Ecsit^+/+^* animals. Following stimulation with lipopolysaccharide (LPS), the levels and phosphorylation of p38-MAPK and JNK were determined by western blot. Results show no significant alterations in the TLR response to stimulation with LPS between Ecsit^+/+^ and *Ecsit^N209I/N209I^* BMDMs, indicating that the N209I mutation of the ECSIT protein does not significantly affect the function of this protein in signal transduction through the TLR pathway (**FigFig EV5 B-C**).

Finally, the interaction between ECSIT and TRAF6 protein was determined by co-immunoprecipitation. 6xHis tagged ECSIT (pCMV6-AC-HIS-ECSIT or pCMV6-AC-HIS-ECSIT^N209I^) and Myc tagged TRAF6 (pCMV6-Entry-TRAF6) were overexpressed in HEK293T cells and cell lysate collected for co-immunoprecipitation with antibodies against the relevant tags. Results (**Fig 3 B**) confirm the interaction between both 50kDa and 45kDa wild-type ECSIT and 60kDA TRAF6 proteins. Furthermore, this interaction is maintained with the presence of the N209I mutation in the ECSIT protein, indicating that the N209I mutation does not deleteriously affect the function of the ECSIT protein in this protein-protein interaction.

Taken together, these data indicate that the TLR function of ECSIT protein is not significantly affected by the mutation of ECSIT, however, the assembly of the electron transport chain supercomplex I is significantly impacted by this mutation, and that this effect is most profound in the heart tissue.

### A cardiac mitochondrial defect

Mitochondrial structure was assessed by TEM (**Fig 4 A-F**) and showed that, whilst there was no swelling of mitochondria or significant changes in mitochondrial size, in *Ecsit^N209I/N209I^* animals, there were signs of hypercondensed and disorganised christae in both interfibrillar and perinuclear mitochondria, suggestive of mitochondrial dysfunction. Western blot analysis of the inner and outer mitochondrial membrane proteins COXIV and TOMM20 confirmed that there were no differences in mitochondrial protein levels between *Ecsit^N209I/N209I^* hearts and wild type (**Fig 4 G-H**). However, the master regulator of mitochondrial biogenesis, PGC1α, was significantly elevated (57%, p=0.0016) in *Ecsit^N209I/N209I^* hearts (**Fig 4 I**), indicating that there is an upregulation of the mitochondrial biogenesis pathway.

**Fig 4:**
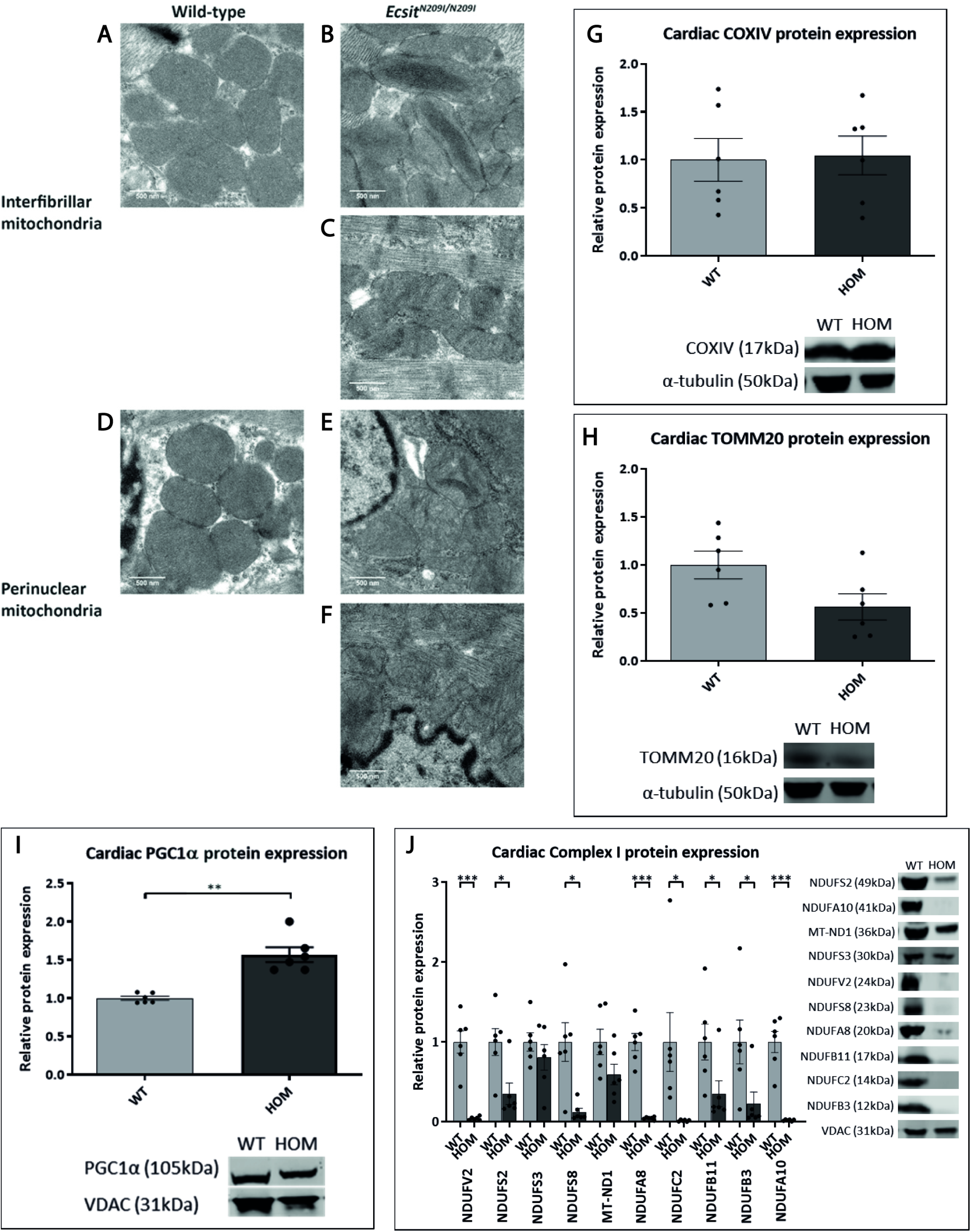
Representative TEM images of wild-type and *Ecsit^N209I/N209I^* interfibrillar and perinuclear mitochondria from cardiac tissue demonstrating structural abnormalities observed in EcsitN209I/N209I samples. Wild-type interfibrillar (A) and perinuclear (B) mitochondria demonstrate consistent evenly stacked cristae with no signs of disorganisation. Mutant interfibrillar mitochondria demonstrate phenotypes such as hyper condensed (C) and disorganised cristae (D). Perinuclear mitochondria from mutant hearts also demonstrate disorganisation (E and F) although hyper-packed cristae were not observed. Scale bars = 500nm. (G-I) Normalised cardiac COXIV, TOMM20 and PGC1α protein expression demonstrating no differences in inner or outer membrane protein abundance but an elevation of mitochondrial biogenesis pathways. (J) Cardiac complex I protein abundance of various proteins representing complex I subunits and accessory proteins, N (V2), Q (S2, S3, S8), PP (ND1, A8, C2), PD (B11, B3) and the accessory protein NDUFA10. Quantification (Normalised to VDAC, relative to WT average) shows a significant reduction in protein levels of all except 2 proteins, NDUFS3 and MT-ND1. Mean ± SEM, *p<0.05, ***p<0.001.

Complex I is the largest of the respiratory chain complexes with distinct domains, each with a dedicated function and sub-assembly process and associated assembly factors. Western blot (**Fig 4 J**) of proteins from each of these subassemblies (N – NDUFV2, Q – NDUFS2, NDUFS3 and NDUFS8, P_p_ – NDUFC2, NDUFA8 and MT-ND1, P_D_ – NDUFB11 and NDUFB3) as well as accessory subunit (NDUFA10) demonstrate a reduction of protein in each of the subunits as result of the N209I mutation in ECSIT. ECSIT was previously demonstrated to be involved early in the assembly of the P_P_ arm of complex I as part of the MCIA complex (Giachin *et al*., 2016; Heide *et al*., 2012) and these results confirm that loss of this essential step has a downstream effect on the continued assembly of the complex.

### Tissue specificity of the mitochondrial defect

To investigate the underlying cause of why no other severe phenotypes were observed as part of the normal screening process (Potter *et al*., 2016), western blots of various complex I proteins were also performed in brain tissue (**Fig 5 A**). These results show a significant reduction several complex I proteins, although less severe than in heart lysate. In particular, complex I protein levels were reduced in brain (~30%), kidney (~30%), and liver (~60%), and not significantly altered in skeletal muscle (**FigFig EV4**). In gel activity assays (**Fig 5 B**) confirmed a reduction in activity of complex I in isolated mitochondria from heart muscle whilst no observable difference in activity levels could be seen in mitochondria isolated from brain. This suggests that the reduction seen in complex I protein levels in brain was insufficient to reduce the activity of the complete enzyme complex. These results were verified by seahorse extracellular flux assay performed on isolated mitochondria from heart and brain tissue of wild-type and *Ecsit^N209I/N209I^* animals (**Fig 5 C**). This confirmed a significant reduction in state III and state IIIu oxygen consumption levels in isolated heart mitochondria from *Ecsit^N209I/N209I^* animals with no changes seen in isolated brain mitochondria from the same animals.

**Fig 5:**
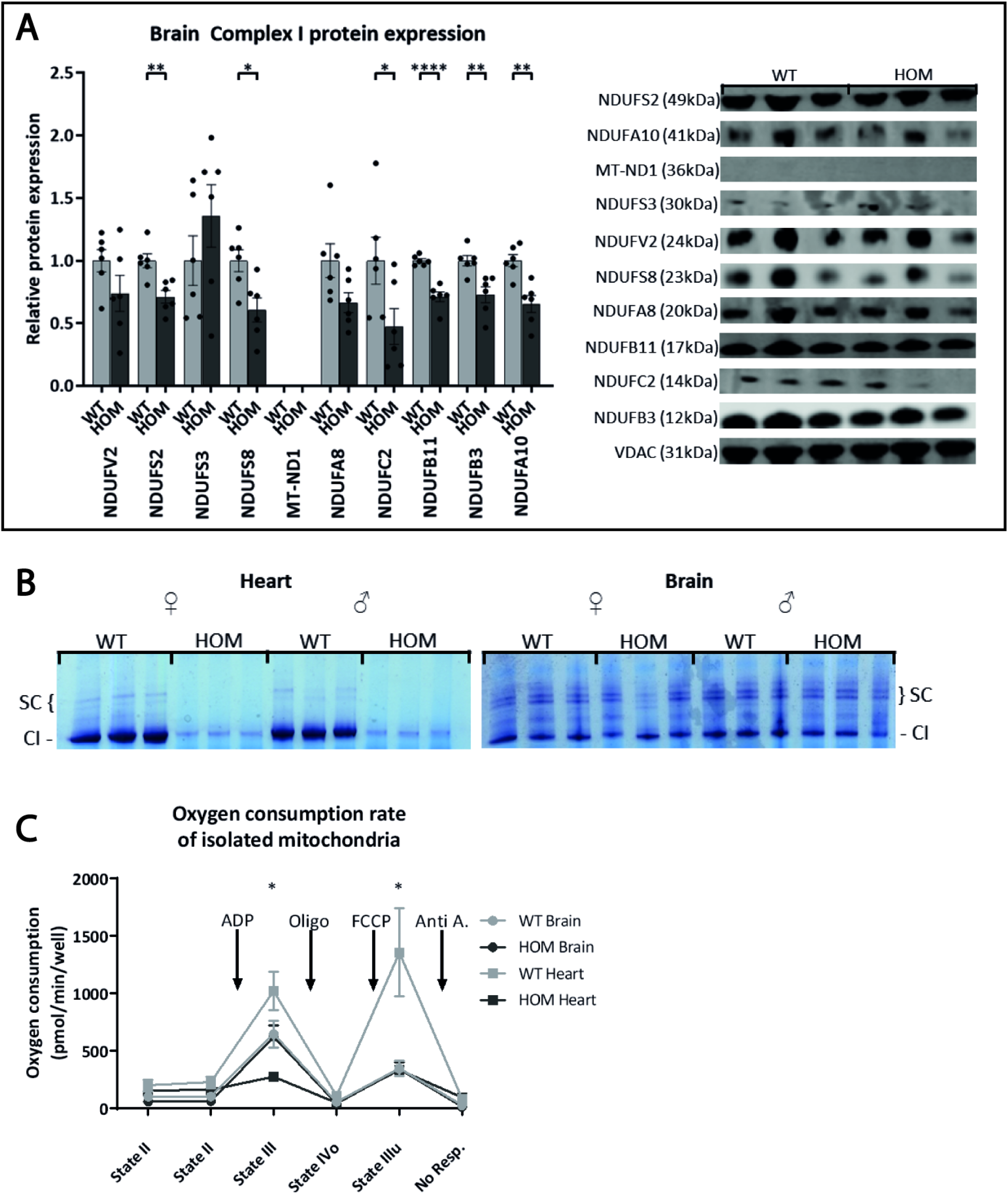
(A) Brain complex I protein abundance of various proteins representing complex I subunits and accessory proteins, N (V2), Q (S2, S3, S8), PP (ND1, A8, C2), PD (B11, B3) and the accessory protein NDUFA10. Quantification (Normalised to VDAC, relative to WT average) shows a significant reduction in protein levels of all except 4 proteins, NDUFV2, NDUFS3, MT-ND1 and NDUFA8. MT-ND1 was not detectable by western blot in brain samples. (B) In gel activity assay of complex I in mitochondria isolated from heart and brain tissue of wild-type and *Ecsit^N209I/N209I^* animals. Depth of stain corresponds to complex I activity with deeper staining reflecting greater activity. Results show a reduction in complex I activity of *Ecsit^N209I/N209I^* hearts, whilst brains show no differences between genotypes. (C) Seahorse oxygen consumption rate measurements from wild-type and *Ecsit^N209I/N209I^* heart and brain mitochondria. Significant differences can be seen between wild-type and *Ecsit^N209I/N209I^* heart mitochondria during state III and state IIIu respiration. No differences are seen between wild-type and *Ecsit^N209I/N209I^* brain mitochondria. Mean ± SEM, *p<0.05, ***p<0.001.

Taken together these data suggest that whilst ECSIT is ubiquitously expressed, its role in complex I assembly differs between tissues, resulting in a less severe mitochondrial deficiency in certain tissues.

### ECSIT protein levels and interactions

Previous work has demonstrated that ECSIT has 2 main isoforms, a 50kDa cytosolic isoform and a 45kDa mitochondrial isoform that is formed following the cleavage of a 5kDa mitochondrial targeting sequence at the proteins N terminus upon localisation to the mitochondria (Vogel *et al*., 2007). Western blot analysis demonstrated the presence of both isoforms in heart and brain tissue (**Fig 6 A-B**), with a significant increase in both protein products in the heart tissue of mutants compared to wild type, without measurable differences in brain tissue. Interestingly, a third band, corresponding to a previously undescribed ~16kDa fragment, can be seen in the heart tissue of wild-type animals, this fragment is conspicuously absent from the heart tissue of *Ecsit^N209I/N209I^* animals as well as in brain liver and kidney (**FigFig EV6**) from both genotypes.

**Fig 6:**
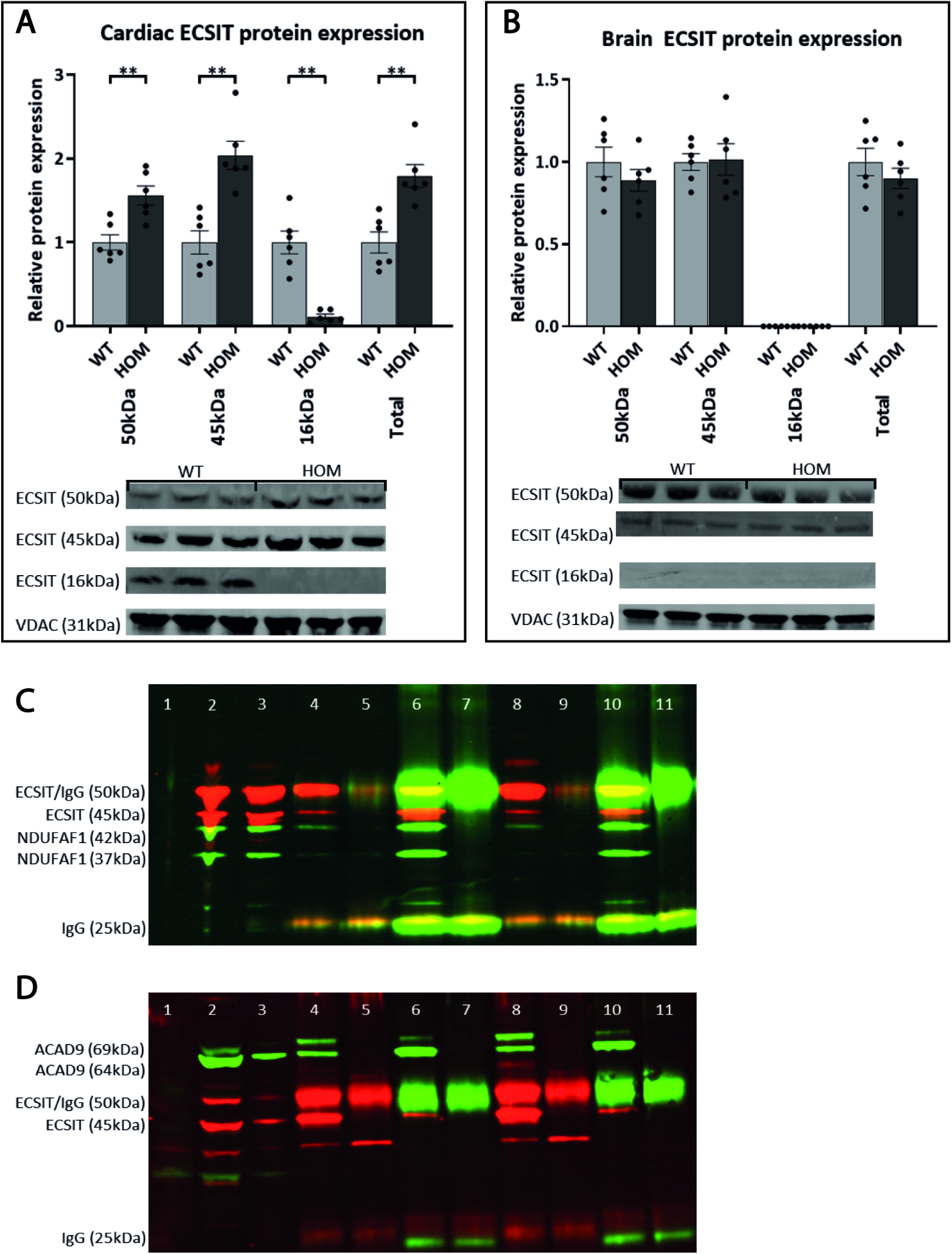
(A) and (B) Cardiac and brain ECSIT protein abundance normalised to loading control VDAC. Results show an increase in both 50kDa and 45kDa ECSIT protein in *Ecsit^N209I/N209I^* animals compared to wild types. In contrast, 16kDa ECSIT demonstrates a significant reduction in *Ecsit^N209I/N209I^* animals, to the point where it is essentially undetectable in cardiac tissue from these animals. (C) and (D) Immunoprecipitation of wild-type and mutant ECSIT (His tagged) (45 and 50kDa) with full length wildtype NDUFAF1 or ACAD9 (Myc tagged) (60kDa) (green – Myc, red – His) 1. Untransfected input lysate, 2. Wild-type ECSIT(His) + wild-type NDUFAF1/ACAD9(Myc) input lysate 3. ECSIT N209I(His) + wild-type NDUFAF1/ACAD9(Myc) input lysate, 4. Wild-type ECSIT(His) + wild-type NDUFAF1/ACAD9(Myc) anti-His immunoprecipitation, 5. Empty AC-His vector + wild-type NDUFAF1/ACAD9(Myc) anti-His immunoprecipitation, 6. Wild-type ECSIT(His) + wild-type NDUFAF1/ACAD9(Myc) anti-Myc immunoprecipitation, 7. Wild-type ECSIT(His) + empty entry(Myc) vector anti-Myc immunoprecipitation, 8. N209I ECSIT(His) + wild-type NDUFAF1/ACAD9(Myc) anti-His immunoprecipitation, 9. Empty AC-His vector + wild-type NDUFAF1/ACAD9(Myc) anti-His immunoprecipitation, 10. N209I ECSIT(His) + wildtype NDUFAF1/ACAD9(Myc) anti-Myc immunoprecipitation, 11. N209I ECSIT + empty entry(Myc) vector anti-Myc immunoprecipitation. Mean ± SEM, **p<0.01, ***p<0.001, ****p<0.0001.

To further investigate the role of ECSIT in complex I assembly and the role of the N209I mutation in its function, the interaction with known binding partners was assessed by co-immunoprecipitation. Tagged ECSIT (His) and either ACAD9 (Myc) or NDUFAF1 (Myc) were co transfected into HEK293T cells. Immunoprecipitation was performed using antibodies against the relevant tags and western blot against the various isoforms performed.

As expected, immunoprecipitation with either anti-his or anti-myc antibody was able to pull down both the wild-type ECSIT and NDUFAF1 proteins in both their cytosolic and mitochondrial isoforms (**Fig 6 C**). The mutant N209I protein was also co-immunoprecipitated with ACAD9 and NDUFAF1.

Similarly, immunoprecipitation with ECSIT and ACAD9 (**Fig 6 D**) demonstrated that both proteins can be pulled down using both the anti-his and anti-myc antibodies that correspond to the relevant tagged proteins. The N209I mutation does not seem to significantly affect this protein:protein interaction as the same results are demonstrated in lanes containing only N209I mutant ECSIT protein).

### Complex I assembly processes in various tissues

Finally, to investigate how the assembly process of complex I is altered in different tissues given the differences seen in complex I protein abundance and activity, two-dimensional blue native PAGE was performed. Proteins representing each of the different portions of complex I were chosen and the patterning of each determined in wild-type and *Ecsit^N209I/N209I^* heart tissue and brain tissue (**Fig 7 A**). NDUFV2 (N), NDUFC2 (P_P_) and NDUFB1 (P_D_-b) all demonstrated a markedly altered abundance in the portion of the band corresponding to intact complex I in mutant animals, whilst NDUFB11 (P_D_-a) showed an accumulation at a band corresponding to an assembly intermediate or isolated protein in *Ecsit^N209I/N209I^* heart tissue. There were no notable changes to proteins in the remaining subcomplexes. The proteins that demonstrated changes in heart tissue were then tested in brain tissue and, with the exception of NDUFB1, which was undetectable by western blot, demonstrated no significant alterations in abundance or patterning between wild-type and mutant tissue.

**Fig 7:**
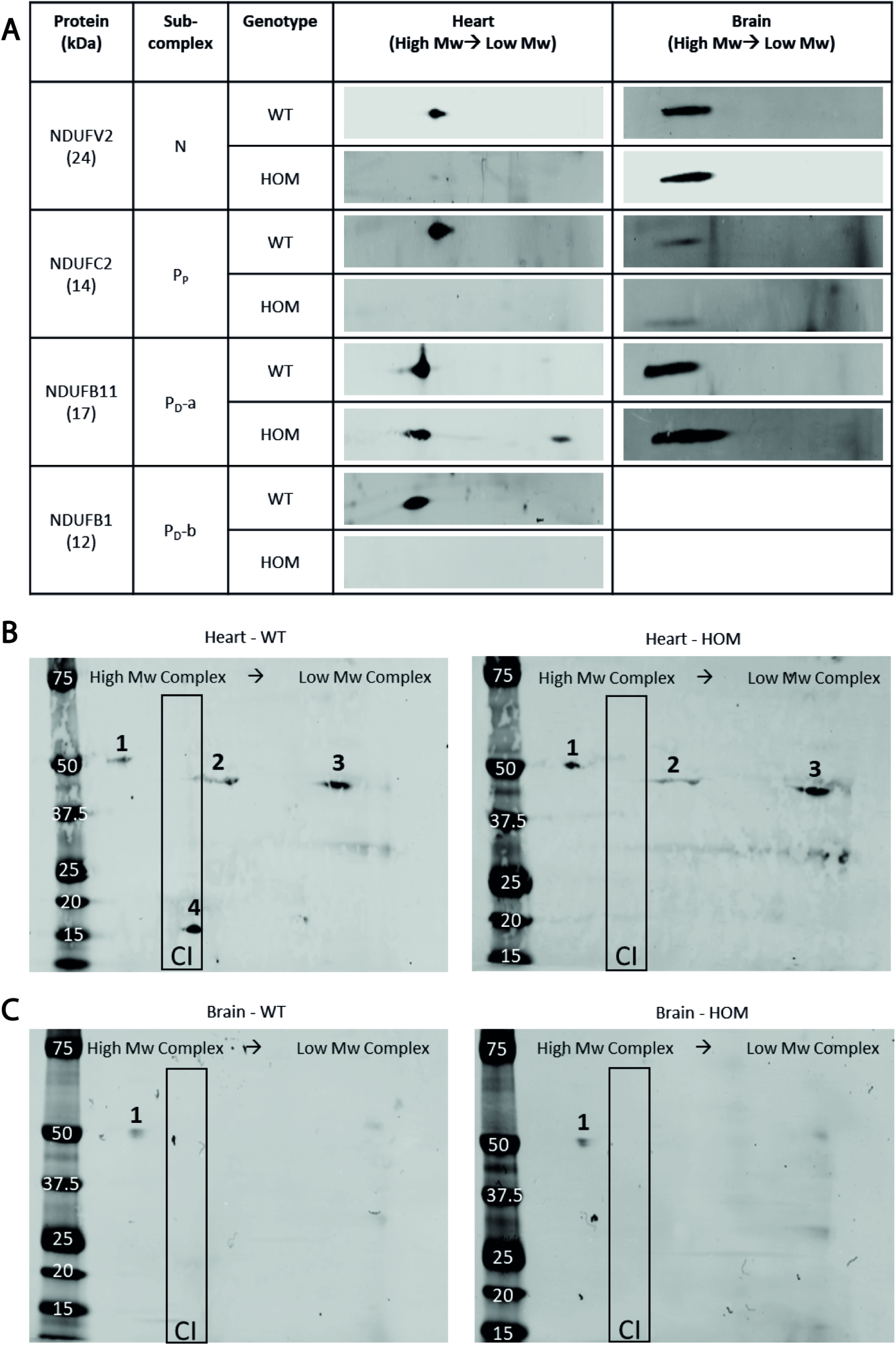
(A) representative 2^nd^-dimensional BN-PAGE blots comparing heart and brain complexes. In each panel high molecular weight complexes are on the left and low weight complexes on the right. Differences seen in the patterning and assembly of complex I sub-complexes in the heart were not seen in the same subcomplexes in the brain. (B) and (C) 2^nd^-dimensional BN-PAGE blots for ECSIT protein in heart and brain demonstrating 50kDa (1) and 45kDa (2 and 3) protein fractions as well as the newly identified 16kDa fragment (4). Blots in brain mitochondria show an absence of both the 45kDa and 16kDa protein fractions. The outlined area is the expected position for intact complex I enzyme associated proteins.

Two-dimensional blue native poly acrylamide gel electrophoresis (2D-BNPAGE) was also performed against ECSIT protein to determine how the N209I mutation affects the assembly process of complex I. Wild-type ECSIT protein (**Fig 7 B**) can be seen in 4 individual bands on the 2D-BNPAGE. The first represents the 50kDA cytosolic isoform and can be seen in a high Mw complex, larger even than fully assembled complex I, which we hypothesise to be the import machinery associated with the outer mitochondrial membrane. The 45kDa mitochondrial fragment can be observed in two bands which correspond to complex I assembly intermediates but is not present in a band that would represent completed complex I. Finally, the 16kDa fragment we had previously identified was present at a size approximately equal to completed, intact complex I holoenzyme. The pattern of the 50 and 45kDa fragments of ECSIT are repeated in the 2D BN-PAGE of the *Ecsit^N209I/N209I^* heart mitochondria, suggesting that the production, and import into the mitochondria, of the mutant ECSIT protein is intact. However, the lack of the 16kDa fragment is striking given the presence of intact complex I in the blots of many of the complex I proteins shown previously. The absence of this fragment in fully assembled complex I suggests that the N209I mutation interferes with the production of this fragment during the assembly process.

2D BN-PAGE in brain mitochondria demonstrate the presence of the 50kDa fragment in a single band (**Fig 7 C**) as seen in the heart mitochondria. As expected, we were unable to detect the 16kDa fragment by western blot in brain lysate (**FigFig EV7**) and in addition, the 45kDa protein product was not detectable by 2D BN-PAGE.

Taken together these data suggests that ECSIT is an essential component of mitochondrial complex I assembly in heart tissue whilst its role in the brain seems not to be essential as both the 45 and 16kDa fragments were absent from the complex I assembly process and in the complete holoenzyme. Furthermore, it suggests that ECSIT is not only involved in the assembly intermediates of complex I in heart mitochondria, but a portion of the protein forms an integral part of completed holoenzyme.

## Discussion

Through a phenotype-driven screen we have identified a point mutation in the mitochondrial complex I assembly factor resulting in tissue specific effects on complex I assembly, primarily affecting cardiac tissue, and resulting in progressive HCM. Mitochondria from *Ecsit^N209I/N209I^* hearts exhibited structural abnormalities, with the presence of hyper condensed, disorganised cristae as well as a small reduction in mitochondrial cross-sectional area of both interfibrillar and perinuclear mitochondria. Protein and mtDNA levels did not demonstrate a change in the overall abundance of mitochondria in heart tissue although the level of PGC1α, the master regulator of mitochondrial biogenesis, was elevated in heart tissue (~50%). This result may indicate that whilst there is an upregulation of the pathway that drives mitochondrial biogenesis, the mutation of ECSIT either inhibits the actual process or results in the production of defective mitochondria, or mitochondrial subunits, that are eliminated by either mitophagy or the mitochondrial unfolded protein response.

Analysis of complex I protein levels confirmed that there was a drastic reduction in complex I levels in heart whereas other tissues (brain, kidney, liver and skeletal muscle) exhibited a smaller reduction. The extent of reduction varies considerably between tissues with the brain showing only a small reduction of around 30% whilst the heart shows a >95% reduction in NDUFB8 protein levels. This reduction in complex I protein level is not reflected across all proteins of complex I, with a more robust reduction in membrane arm proteins than in matrix arm proteins.

Results also demonstrated that, whilst complex I protein levels are reduced in several tissues, the activity of complex I is only affected in cardiac tissue. Comparing heart and brain (two tissues commonly affected by complex I deficiency) using in-gel activity and seahorse analysis of isolated cardiac mitochondria, revealed that complex I activity and mitochondrial respiration were significantly reduced in mutant tissue. However, this was not true for complex I or mitochondria isolated from brain tissue, which showed comparable levels in wild type and *Ecsit^N209I/N209I^* mitochondria in both assays. Similarly, the *Ecsit^N209I/N209I^* mitochondria had little effect on complex I assembly in muscle and other tissues. These data indicate that there are tissue specific differences in the complex I assembly process, and in particular the stage(s) controlled by ECSIT. However, the mutation of ECSIT protein had no impact on the interaction between ECSIT and other proteins of the MCIA complex such as NDUFAF1 and ACAD9, which also showed no changes in protein levels in any tissues tested.

The tissue specific effects of the *Ecsit^N209I/N209I^* mutation explain why the main phenotype observed was HCM. Of interest was the identification of a previously undescribed ~16kDa fragment in wild-type heart tissue, which was absent from *Ecsit^N209I/N209I^* hearts. ECSIT exists as a cytosolic (50kDa) and mitochondrial (45kDa) form in mouse tissues and the levels of these two protein products were higher in *Ecsit^N209I/N209I^* animals compared to wild type. The reason for this accumulation is unclear but the absence of the smaller 16 kDa ECSIT protein in cardiac tissue suggests a cleavage of ECSIT in normal tissue to facilitate the assembly of complex I. It is conceivable that this fragment is a portion of ECSIT produced by the cleavage of the 45kDa ECSIT as part of the normal assembly process for complex I. If this is the case then it may be that the N209I mutation inhibits this cleavage, leading to an accumulation of the larger 45 and 50kDa ECSIT isoforms. In brain tissue, we only observed the 50kDa form of ECSIT in BN-PAGE gels. This may be because ECSIT is not transported into the mitochondria in the brain. This fits with our observations that ECSIT is not required for mitochondrial complex I assembly in brain tissue.

To assess complex I assembly, first and second dimensional blue-native PAGE were performed on mitochondria extracts from heart and brain of wild-type and *Ecsit^N209I/N209I^* animals. This revealed that some aspects of complex I assembly were defective in cardiac tissue from *Ecsit^N209I/N209I^* animals, but not in brain tissue. These defects mainly relate to the assembly of the membrane arm of complex I although some loss of other subunits was also seen. Interestingly, none of the same defects were seen in mitochondria isolated from brain tissue of the same animals. This further supports the hypothesis that complex I assembly is not a universal process but has tissue specific intricacies.

Taken together, these data suggest that the N209I mutation of ECSIT has no effect on protein synthesis, as indicated by the presence of the 50 and 45kDa bands, or on the interaction between ECSIT and the MCIA proteins NDUFAF1 and ACAD9. Our novel finding was the presence of a 16kDa fragment detected by the anti-ECSIT mAb. This fragment was not detected in mutant cardiac tissue or in wild-type and mutant brain tissue. Complex I assembly and activity in brain tissue was unaffected and those tissues tested that did show complex I protein level differences are less severely affected than heart tissue.

In wild-type animals ECSIT is expressed as full length 50kDa protein in all tissues and according to demand is poly-ubiquitinated and targeted to mitochondria where it is imported and the mitochondrial targeting sequence cleaved, leaving a 45kDa ECSIT protein. Once localised to the mitochondria, ECSIT forms part of the MCIA complex along with NDUFAF1 and ACAD9 (Giachin *et al*., 2016). To this point in the complex I assembly our data suggests that ECSIT functions normally. The MCIA complex is then involved in the assembly of the membrane arm of complex I (Heide *et al*., 2012; Nouws *et al*., 2010; Vogel *et al*., 2007). From this point, the mutant ECSIT protein prevents the normal assembly of complex I in cardiac tissue.

However, the assembly process appears to be tissue specific; the N209I mutation varies in its effect on complex I assembly in different tissues and there are even differences in the ECSIT protein association with complex I between brain and cardiac tissue in wild-type animals. The different effects of the N209I mutation on complex I assembly between tissues, possibly because of tissue specific requirements for ECSIT in this process.

In summary, we have identified a novel model of HCM resulting from a mutation in the mitochondrial complex I assembly protein, ECSIT. The mutation has no effect on the TLR pathway but appears to affect the cleavage of ECSIT during mitochondrial complex I assembly. We also provide evidence for tissue specific requirements for ECSIT in complex I assembly, which explains the severe cardiac phenotype in the absence of other phenotypes and provides the first evidence of a mechanism underlying a tissue specific effect of mitochondrial dysfunction. This mutant line provides opportunities to investigate the mechanisms underlying not only HCM but also tissue specific differences in complex I assembly. Furthermore, these findings also have implications for the clinical diagnosis of cardiac disease and testing procedures for complex I assembly deficiencies as this is often carried out using skin fibroblasts (Baertling *et al*, 2017; Koopman *et al*, 2008; Verkaart *et al*, 2007), which may not reflect complex I assembly in all tissues.

## Methods and Materials

### Mice

C57BL/6J and C3H-C3pde6b+ inbred mice were maintained in the Mary Lyon Centre in Harwell UK, in specific pathogen-free conditions. All animal procedures were carried out under the guidance issued by the Medical Research Council in “Responsibility in the Use of Animals for Medical Research” (July 1993) and in accordance with Home Office regulations (Home Office Project Licence No. 30/3070)

### Generation of Mutagenized Mice

The original mutant mouse pedigree was derived from a G_3_ pedigree produced in the MRC Harwell ENU mutagenesis screen as described previously (Potter *et al*., 2016). Briefly, male C57BL/6J mice were mutagenized with ENU and then mated to female C3H.Pde6b+ mice to generate G_1_ founder males, heterozygous for ENU induced mutations. G_1_ males were subsequently bred to female C3H.Pde6b+ mice to generate G_2_ offspring. Lastly, G_2_ females were mated back to the original G_1_ founder to generate two G_3_ cohorts, both heterozygous and homozygous for ENU induced mutations.

### Mapping and Next Generation Sequencing

DNA from affected mice and littermate controls were tested on the Illumina Golden Gate “Mouse MD Linkage Panel” (Oxford Genomics Centre, Wellcome Trust Centre for Human Genetics). DNA from the G_1_ founder of the pedigree was sent for whole genome sequencing (WGS) employing the Illumina HiSeq platform (Oxford Genomics Centre, Wellcome Trust Centre for Human Genetics) and analysed as previously described (Potter *et al*., 2016). The *Ecsit* mutation was validated using Sanger Sequencing (Source Bioscience).

### Light and Electron Microscopy

For light microscopy, hearts fixed in 10% neutral buffered formalin were embedded in paraffin wax and sectioned using a Finesse ME+ microtome (Thermo Fisher). Transverse sections (T/S) were stained with Haematoxylin and Eosin (H&E).

For transmission electron microscopy (TEM), 1mm^3^ cubes of left ventricular tissue were fixed in 3% glutaraldehyde and 4% formaldehyde in 0.1 M PIPES and post-fixed with 1% osmium tetroxide in 0.1 PIPES. Samples were taken from the left ventricular free wall of 3 wild type and 3 *Ecsit^N209I/N209I^* males at 16 weeks of age. After serial dehydration in increasing concentration of ethanol, the tissue was embedded in epoxy resin (TAAB) and polymerised overnight at 60°C. Golden ultrathin sections (70-80 nm) were cut with a diamond knife and collected on copper/palladium grids. To improve contrast, blocks were stained with 2% uranyl acetate and grids were stained with lead citrate.

Images were collected at the Wolfson bioimaging facility at the University of Bristol using a Tecnai 12 Biotwin electron microscope.

### Echocardiography

Echocardiography was performed by the phenotyping core of the Mary Lyon centre at MRC Harwell. Twelve-week-old male and female mice were anaesthetised with 4% isoflurane (maintained at 1.5%) in oxygen and echocardiogram performed using a Vevo 770 high-resolution *in vivo* micro imaging system with a Visualsonics RMV707B Probe (30 MHz). Body temperature was monitored using a rectal thermometer and maintained using a heat lamp at 36-38°C. ECG monitoring was performed using limb electrodes, and the heart rate maintained at or above 400bpm. Short axis B and M mode images were taken using the papillary muscles as a point of reference for positioning of the probe. Image contrast and gain functions were used for clarity and frame rate of 110Hz used throughout. Measurements were taken from M-mode images using the inbuilt Vevo software.

### Western Blot

Proteins were extracted from tissue by homogenising in RIPA buffer (150mM NaCl, 1% NP-40, 0.5% DOC, 0.1% SDS, 50mM Tris, pH 7.5) containing phosphatase (Roche) and protease (Roche) inhibitors in precellys CK28 homogenisation tubes.

Proteins were isolated from macrophages by scraping the macrophages from the plate and centrifuging to obtain the cell pellet before lysing in RIPA buffer containing phosphatase and protease inhibitors.

Protein concentration was measured by Bradford assay (Bio-rad) and measured on a μQuant plate reader (Biotek instruments inc., VT, USA). 20μg of total protein was mixed with LDS sample buffer (Invitrogen) and reducing agent (Invitrogen) loaded onto NuPAGE™ 4-12% Bis-Tris protein gels (Invitrogen). Electrophoresis was performed at 200V for 60 minutes at room temperature in 1x MOPS.

PVDF membrane (GE) was activated in absolute methanol for 1 minute and proteins transferred using X-Cell blot module (Invitrogen) containing 1x transfer buffer (Invitrogen), 20% methanol and 1x antioxidant (Invitrogen) according to manufacturer’s instructions.

Following transfer, membranes are blocked in either 5% w/v milk powder in phosphate buffered saline (PBS) containing 0.1% tween or 5% BSA (bovine serum albumin) in TBS containing 0.1% tween (for phosphorylated proteins) for 60 minutes with shaking. Primary antibodies diluted in 5% milk/PBS-T or 5% BSA/TBS-T are incubated overnight at 4°C before three, 5 minute washes in PBS-T or TBS-T. Secondary antibodies (Section 2.5) are diluted in 5% milk/PBS-T or 5% BSA/TBS-T and incubated for 1 hour. Fluorescent secondary antibodies are protected from light during this and subsequent steps. Membranes are subsequently washed a further 3 times before drying. Blots were scanned using a LI-COR Odyssey Cl-x or SA scanner (LI-COR Biosciences, Cambridge, UK). Image Studio Lite software (LI-COR Biosciences) is used for quantification and analysis (Median, 3 pixel border background).

### Flow cytometry analysis

Immune cell profiling in blood was performed by flow cytometry as described previously (Vikhe *et al*, 2020). Samples were collected from 12-week-old male and female wild-type and *Ecsit^N209I/N209I^* animals. Briefly blood was collected by retro-orbital bleeding in lithium heparin tubes, from the mice under terminal anesthesia induced by an intraperitoneal overdose of sodium pentobarbital. For flow cytometry analysis, 50 μl of blood was resuspended in 1 ml of red blood cell (RBC) lysis buffer (Biolegend™) for 10 min on ice followed by two washes with 1 ml phosphate buffer saline (PBS). The final pellet was resuspended in FACS buffer (5 mM EDTA, 0.5% fetal calf serum in PBS) and transferred to V-bottom 96 well plate. The cell suspension was incubated for 15 min with CD16/CD32 antibody (BD Pharmingen) at a dilution of 1:100. After centrifugation at 800 × g for 1 min, the cell pellets were resuspended in 100 μl of antibody cocktail: F4/80 – PE (1:200) (e biosciences™), CD11b – PE – CF594 (1:200), Ly6G – BV421 (1:200), Ly6C – FITC (1:200) and CD5 – BD421 (1:800) (BD Horizon™) and incubated for 20 min in the dark. Cells were centrifuged and washed twice with FACS buffer and finally fixed by adding 210ul of 0.5% PFA. Flow cytometry was performed using a BD FACSCanto II system and FlowJo software (Tree Star) was used to analyze the data obtained.

### Macrophage Culture

12-week-old wild-type and *Ecsit^N209I/N209I^* animals were sacrificed by cervical dislocation and bone marrow flushed from femur and tibia with PBS containing 0.6mM EDTA. Cell suspension was filtered through a pre-wetted 70μm cell strainer. Cells were pelleted at 400xg for 7 minutes (4°C). Red blood cells were removed by resuspending the pellet in 3mls of red blood cell lysis buffer (155mM NH_4_Cl, 12mM NaHCO_3_, 0.1mM EDTA) and incubated for 1 minute before diluting in 10mls of PBS (0.6mM EDTA) and again pelleted at 400xg.

Finally, cells were plated at a concentration of 2.5×10^6^ cells/mL in DMEM (pyruvate, glutamine) containing 10% FBS, 100U/ml penicillin-streptomycin and 100ng/ml of macrophage colony stimulating factor (MCSF) (Cell Guidance Systems). Cells were maintained at 37°C, 5% CO_2_ with media changes on day 3 and 6. On day 7 cells were harvested by manual scraping with PBS (0.6mM EDTA). Cells were counted again and re-plated at a concentration of 6×10^6^ cells/well of a 6-well plate in DMEM (pyruvate, glutamine), 10% FBS, 100U/ml penicillin-streptomycin, 100ng/mL MCSF. Plated cells were activated with 100ng/ml of lipopolysaccharide (LPS) added directly to the media and incubated for 24 hours at 37°C, 5% CO_2_.

Activated cells were harvested by manual scraping with PBS (0.6mM EDTA).

### Cloning and Transfection

Full length *Ecsit* clone (Dharmacon) was ligated into pCMV6-AC-HIS vector following incorporation of flanking restriction enzyme sites by PCR (SgfI AGGCGATCGCCATGAGCTGGGTGCAGGTCAACTT, MluI – GCGACGCGTACTTTGCCCCTGCTGCTGCTCTG) and grown in XL-10 Gold E.Coli (Agilent). Site directed mutagenesis (Q5-SDM – NEB) was utilized to introduce the N209I mutation (Forward – CGATTCAAGATTATCAACCCCTAC, Reverse – GGTGAACCACAGCTTCATC). Full length Ndufaf1 (Dharmacon MMM1013-202762755) was similarly incorporated into pCMV6-Entry (SgfI – GAGGCGATCGCCATGTCTTCCATTCACAAATTACT, MluI – GCGACGCGTTCTGAAGAGTCTTGGGTTAAGAA). Vector DNA isolated by QIAprep Spin Miniprep kit (Qiagen).

pCMV6-AC-HIS-Ecsit (wild-type or *Ecsit^N202I^*), pCMV6-Entry-TRAF6, pCMV6-Entry-ACAD9, pCMV6-Entry-NDUFAF1 were transfected into Hek293T cells using jetPRIME reagent (Polyplus). Hek293T cells were plated at 2.5×10^5^ cells/well (6 well plate) in DMEM (high glucose, glutamax, 10% FBS, 100U/ml penicillin streptomycin).

### Co-IP

48 hours following transfection, HEK293T cells transfected with relevant vectors to express ECSIT or associated proteins were briefly washed with PBS and lysed in RIPA buffer (150mM NaCl, 1% NP-40, 0.5% DOC, 0.1% SDS, 50mM Tris, pH 7.5) with protease inhibitors (Roche). Protein concentration was assessed by Bradford assay (Bio-Rad) and protein diluted to 1mg/ml 1ml of protein lysate was precleared with 20μl of protein G sepharose bead slurry (Sigma) for 1 hour to remove native immunoglobulins. Protein G beads were removed by briefly spinning at 1000xg for 1 minute and the supernatant incubated with 4μg of relevant antibody (outlined in key resources table) over night to bind the protein of interest. Following antibody binding, lysate was incubated with 20μl of protein G sepharose bead slurry for 1 hour to bind antibody and attached protein/s. Beads were again pelleted by centrifugation at 100xg for 1 minute and washed 3 times in RIPA buffer. Beads were left in 30μl of RIPA buffer and 1x LDS sample buffer (Invitrogen) and reducing agent (Invitrogen) added before boiling the sample at 95°C for 10 minutes to dissociate the beads from the bound antibody.

Samples were loaded onto NuPAGE™ 4-12% Bis-Tris protein gels (Invitrogen) and run as with western blots.

### Mitochondrial Isolation

Mitochondria were isolated from frozen heart and brain tissue taken from wild-type and *Ecsit^N209I/N209I^* animals. Hearts were lysed in 10ml/g of homogenisation medium A (0.32M sucrose, 1mM EDTA, 10mM tris-HCl, pH7.4, filter sterilised) in an Elvehjem-Potter homogeniser on ice by hand. Homogenate was centrifuged at 1000xg or 5 minutes at 4°C and supernatant retained. Supernatant was centrifuged for 2 minutes at 9000xg at 4°C before removing the supernatant and fluffy coat, leaving behind the mitochondrial pellet. The mitochondrial pellet was subsequently resuspended in 100μL of homogenisation medium A and centrifuged at 9000xg for 10 minutes at 4°C, discarding the supernatant. This wash step is repeated 5 times and the mitochondrial pellet stored at −80°C. Brains were homogenised in the same fashion using 5ml/g of homogenisation medium AT (0.075M sucrose, 0.225M mannitol, 1mM EGTA, 10mM tris-HCl, pH7.4, filter sterilised). Mitochondria were resuspended in 100uL of homogenisation medium and the concentration determined by Bradford assay (Bio-rad).

### In Gel Activity

50μg of isolated mitochondria was resuspended in 1x native sample buffer (Invitrogen) 2% digitonin and 1x protease inhibitors (Roche) and incubated for 1 hour on ice before addition of 0.5% G-250 sample additive (Invitrogen). Prepared samples were run on NativePAGE™ 3-12% Bis-Tris gels under native conditions at 150V for 30 minutes with 1x native cathode buffer (Invitrogen) followed by 90 minutes at 250V in 0.1x native cathode buffer. The gel was then incubated for 1 hour in 150μM NADH, 3mM nitro blue tetrazolium and 2mM Tris-HCl (pH 7.4). Activity of Complex I corresponds to depth of blue-purple stain.

### Seahorse

Mitochondria for seahorse analysis were isolated from freshly dissected hearts and brains obtained from wild-type and *Ecsit^N209I/N209I^* animals. Samples were kept on ice and homogenised in MSHE+BSA (70mM sucrose, 210mM Mannitol, 5mM HEPES, 1mM EGTA, 0.2% w/v FFA free BSA, 1M KOH) in a Dounce homogeniser using both A and B pestles. Mitochondria were separated from lysate by subsequent centrifugation 800xg for 10 minutes at 4°C twice, discarding the pellet after each step. The supernatant is retained and centrifuged at 8000xg for 10 minutes at 4°C. The remaining mitochondrial pellet is suspended in MSHE+BSA and the concentration measured using a Bradford assay (Bio-rad).Isolated mitochondria are diluted 1:10 in ice cold mitochondrial assay solution (MAS) (70mM sucrose, 220mM mannitol, 10mM KH_2_PO_4_, 5mM MgCl_2_, 2mM HEPES, 1mM EGTA, 0.2% w/v FFA free BSA, 1M KOH) before diluting further in MAS to the desired concentration and loaded onto Seahorse XF24 plates at a concentration of 5μg/well. The plate is centrifuged at 2000xg for 20 minutes at 4°C to ensure mitochondria are adherent.

450μL of MAS containing 10mM pyruvate and 10mM malate is added to the mitochondria in the plate and the plate incubated at 37°C for 10 minutes without CO_2_. Measurements are made on the seahorse with the addition of 40mM ADP (4mM final), 20μM Oligomycin (2μM final), 40μM FCCP and Antimycin A (4μM final). Measurements were taken over a period 3 minutes flanked by 1-minute mix periods.

### BN-PAGE

250ug of isolated mitochondria was resuspended in 1x native sample buffer (Invitrogen) 2% digitonin and 1x protease inhibitors (Roche) and incubated for 1 hour on ice, centrifuged at 20000xg for 30 minutes at 4°C before supernatant is removed and addition of 0.5% G-250 sample additive (Invitrogen). First dimensional BN-PAGE was run on NativePAGE™ 3-12% Bis-Tris gels under native conditions at 150V for 30 minutes with 1x native cathode buffer (Invitrogen) followed by 90 minutes at 250V in 0.1x native cathode buffer. Second dimensional BN-PAGE, first dimensional lanes are cut from the gel and incubated in 1% SDS with 1% beta-mercaptoethanol for 1 hour before the gel slice was loaded into an NuPAGE 4-13% 1.0mmx2D gel (Invitrogen). Gels were run at 200V for 50 minutes at room temperature in MOPS. Transfer for PVDF membrane and remaining steps were performed as with Western Blot.

### Muscle Fibretyping

Soleus and extensor digitorum longus (EDL) from male wild-type and *Ecst^N209I/N209I^* animals and frozen over isopropanol on liquid nitrogen. Following freezing, frozen muscle samples were mounted in Tissue Tech freezing medium (Jung) and cooled by dry ice/ethanol. Cryosections taken at 10μM thick were dried for 30 minutes at room temperature before being washed three times in PBS and incubated in permeabilisation buffer solution (4mM HEPES, 3mM MgCl_2_, 10mM NaCl, 1.5mM Sodium azide, 60mM Sucrose, 0.1% Triton X-100) for 15 minutes. Following permeabilisation, samples were washed in wash buffer (1x PBS with 5% foetal calf serum (v/v) and 0.05% Triton X-100) for 30 minutes at room temperature.

Primary antibodies against MHCI, MHCIIA, MCHIIX and MHCIIB were diluted in wash buffer and incubated overnight at 4°C. The next day samples are incubated in secondary antibody diluted in wash buffer for 1 hour in darkness. Finally, slides were mounted in fluorescent mounting medium and myonuclei visualised with 2.5μg/ml DAPI.

Fluorescence microscopy was performed with Zeiss AxioImegar AI an images captured with Axiocam digital camera. Analysis was performed with Zeiss Axiovision computer software version 4.8.

### Statistical Analysis

All statistical analysis was carried out using Microsoft Excel and Graphpad Prism 8 (GraphPad Inc.). All analysis is displayed as mean ± standard error of the mean (SEM). Student’s T-test was used where a single variable accounts for differences in two groups and one-way ANOVA with Tukey’s multiple comparisons test for multiple groups separated by a single variable. Differences were considered significant at p values of less than 0.05. *p<0.05, **p<0.01, ***p<0.001, ****p<0.0001.

## Acknowledgements

The authors would like to thank Professor Sankar Ghosh (Columbia University, New York, USA) for supplying the *Ecsit^-/-^* mouse.

This work was primarily funded by the Medical Research Council Award MC U142684172 with CV funded by Medical Research Council award (MC_UU_00015/5).

## Author Contributions

TN and PKP conceived the work, designed the project and drafted the manuscript. TN, SF, AB, PV, GC, SSO acquired data. CV and KP interpreted data and approved the final version.

## Conflict of Interests

None declared

